# Double-digest RAD sequencing outperforms microsatellite loci at assigning paternity and estimating relatedness: a proof of concept in a highly promiscuous bird

**DOI:** 10.1101/169144

**Authors:** Derrick J. Thrasher, Bronwyn G. Butcher, Leonardo Campagna, Michael S. Webster, Irby J. Lovette

**Affiliations:** Fuller Evolutionary Biology Program, Cornell Laboratory of Ornithology, Ithaca, NY 14850, USA; Department of Ecology and Evolutionary Biology, Cornell University, E145 Corson Hall, Ithaca, NY 14853, USA; Macaulay Library, Cornell Lab of Ornithology, 159 Sapsucker Woods Rd, Ithaca, NY 14850, USA; Department of Neurobiology and Behavior, Cornell University, W361 Mudd Hall, 215 Tower Rd, Ithaca, NY 14853, USA

**Keywords:** double-digest restriction site-associated DNA sequencing (ddRAD-seq), microsatellite, single nucleotide polymorphism (SNP), parentage, relatedness, cooperative breeding

## Abstract

Information on genetic relationships among individuals is essential to many studies of the behavior and ecology of wild organisms. Parentage and relatedness assays based on large numbers of SNP loci hold substantial advantages over the microsatellite markers traditionally used for these purposes. We present a double-digest restriction site-associated DNA sequencing (ddRAD-seq) analysis pipeline that, as such, simultaneously achieves the SNP discovery and genotyping steps and which is optimized to return a statistically powerful set of SNP markers (typically 150-600 after stringent filtering) from large numbers of individuals (up to 240 per run). We explore the tradeoffs inherent in this approach through a set of experiments in a species with a complex social system, the variegated fairy-wren (*Malurus lamberti*), and further validate it in a phylogenetically broad set of other bird species. Through direct comparisons with a parallel dataset from a robust panel of highly variable microsatellite markers, we show that this ddRAD-seq approach results in substantially improved power to discriminate among potential relatives and considerably more precise estimates of relatedness coefficients. The pipeline is designed to be universally applicable to all bird species (and with minor modifications to many other taxa), to be cost- and time-efficient, and to be replicable across independent runs such that genotype data from different study periods can be combined and analyzed as field samples are accumulated.

## Introduction

Advances in molecular techniques over the past several decades have substantially improved our ability to test questions about animal social behavior by providing reliable information on the genetic relationships among individuals (Westneat *et al.* 1990; Hughes 1998; Avise *et al.* 2002; Griffith *et al.* 2002; Solomon *et al.* 2004; Myers & Zamudio 2004). Microsatellites have been the molecular ‘tool-of-choice’ for this application since the 1990s, as microsatellite loci are often highly polymorphic, with up to dozens of co-segregating alleles at a single locus (Queller *et al.* 1993; Li *et al.* 2002; Selkoe & Toonen 2006; Guichoux *et al.* 2011). Accordingly, a small number of highly variable microsatellite loci can provide considerable power for discerning genetic relationships among individuals (Queller *et al.* 1993; Blouin 2003; Webster & Reichart 2005). However, microsatellite assays also have some practical drawbacks.

Microsatellite laboratory protocols developed for one species are often not suitable for use in other species because the primers may not amplify well and targeted loci are often not as polymorphic, especially in more distantly related taxa (Galbusera 2000; Decroocq *et al.* 2003; Hedgecock *et al.* 2004; Primmer *et al.* 2005). Next-generation sequencing has made the discovery of microsatellite loci for individual species more attainable (Davey *et al.* 2011). However, discovering microsatellite loci can be very time consuming and costly, largely due to protracted testing and optimization of candidate primers after the initial sequencing. Additionally, traditional PCR-based microsatellite assays also incur substantial financial and lab-bench time investments. The manual scoring of microsatellite alleles also requires substantial researcher time, and can involve various forms of error arising from alleles that have more than one clearly defined peak, allelic drop-out and null allele issues, and the various sources of human error that are inherent in any complicated workflow (Pemberton *et al.* 1995; Hedgecock *et al.* 2004; Hoffman & Amos 2005; Kalinowski *et al.* 2007).

Many of these limitations are less severe in assays based on single-nucleotide polymorphisms (SNPs), which require fewer steps and have greater automation (Gut 2001; Syvänen 2001; Seeb *et al.* 2011; Davey *et al.* 2011). SNPs are appropriate alternatives for studies of parentage and relatedness data because they are abundant in the genome, have low mutation rates (Brumfield *et al.* 2003; Morin *et al.* 2004), and can be scored semi-automatically (Garvin *et al.* 2010; Guichoux *et al.* 2011). In comparison to microsatellite-based relationship tests, the primary limitation of SNPs is that they are typically biallelic, whereas microsatellite loci are often multiallelic, and hence the statistical power of SNP loci for discriminating parentage and relatedness is far lower on a per-locus basis (Ball *et al.* 2010). Compared to highly variable microsatellite loci, a substantially higher number of SNP markers is therefore required to achieve appropriate power in parentage and relatedness studies (Glaubitz *et al.* 2003; Morin *et al.* 2004; Coates *et al.* 2009).

Recently, the application of SNPs for use in analyses of parentage, relatedness, and overall population structure has received greater attention (Glaubitz *et al.* 2003; Anderson & Garza 2006; Coates *et al.* 2009). Studies in birds (Cramer *et al.* 2011; Weinman *et al.* 2015; Kaiser *et al.* 2017), fish (Hauser *et al.* 2011), and several domesticated taxa (Tokarska *et al.* 2009; Fernández *et al.* 2013) have developed sufficiently large SNP panels to attain a comparable, if not better, level of resolving power as highly polymorphic microsatellite panels. While each of these studies manage to identify powerful SNP panels, the SNP genotyping methods used are often labor intensive, requiring a significant amount of preparatory work at the discovery stage prior to genotyping of large numbers of individuals. Many of these methods also rely on reference genomes (Anderson & Garza 2006; Heylar *et al.* 2011), or other genomic resources (Fernández *et al.* 2013; Weinman *et al.* 2015; Kaiser *et al.* 2017) (e.g. transcriptome, SNP microarray), for SNP identification. Ultimately, this has afforded several beneficial examples of the utility of SNPs for parentage and relatedness analyses, but without an efficient, universal method of SNP discovery and identification.

Restriction site-associated DNA sequencing (RAD-seq) is a reduced-representation genomic technique that is widely used in molecular genetic studies (Davey & Blaxter 2010; Etter *et al.* 2012; Puritz *et al.* 2014), particularly for linkage and quantitative trait locus (QTL) mapping (Baird *et al.* 2008), genome wide association studies (Davey *et al.* 2011), and phylogeography (Andrews *et al.* 2016). RAD-seq uses restriction enzymes to fragment and sample a fraction of a genome; as it identifies SNPs with no prior knowledge of the genome, it provides a more universal method of SNP discovery (Willing *et al.* 2011). Double-digest restriction site-associated DNA sequencing (ddRAD-seq) is a RAD-seq protocol that allows for selection of an even smaller fraction of the genome through a size selection step, affording the ability to target a smaller total number of SNPs in a greater number of individuals (Peterson *et al.* 2012; Puritz *et al.* 2014; Kess *et al.* 2016). This ability, in concert with the fact that no prior knowledge of the genome is needed, makes ddRAD-seq an attractive method of simultaneous SNP discovery and screening for use in discerning genetic relationships among individuals.

Here, we describe a ddRAD-based approach to the simultaneous discovery and screening of high numbers of SNP loci with high power for testing questions about parentage and relatedness. These protocols are optimized to generate an appropriately robust set of SNP markers for 240 individuals per run, to be repeatable across runs to allow the combination of SNP datasets generated at different times, and to be universally applicable to birds (and with small modifications, to other organisms) without requiring a species-specific marker discovery step. We validate these methods by conducting a SNP-based parentage and relatedness study in the highly promiscuous, and socially complex, variegated fairy-wren *(Malurus lamberti).* We compare the results with previously generated paternity assignments and relatedness information, based on microsatellite screens of the same fairy-wren individuals and social groups. To illustrate the broad utility of this method we report the number of loci recovered for equivalent studies of parentage that included different numbers of individuals (from less than 10 to almost 500) of a variety of other species that collectively span much of the phylogenetic diversity of living birds.

## Methods & Materials

### Study population

The variegated fairy-wren, endemic to Australia, is a cooperatively breeding bird that lives in social groups composed of kin and non-kin (Schodde 1982; Rowley & Russell 1997). Male dispersal is limited, and rates of extra-pair fertilizations (EPFs) are high (~68% of all young, assessed with a panel of 12 species-specific microsatellites; DJ Thrasher, unpublished data). We intensively monitored a color-banded population of the nominate subspecies, *M. l. lamberti,* on Lake Samsonvale (27°16’S, 152° 41’E), 30 km northwest of Brisbane, Queensland, Australia, from 2012 – 2016. The population ranges from about 250-300 adults depending on year-to-year conditions. The study site is bounded on most sides by Lake Samsonvale, and on its westernmost side by a major highway, which increases our confidence in sampling most, if not all, of the adults in the population. We also monitored all nesting attempts to measure, mark, and collect blood samples from nestlings 6 days after hatching. Blood samples were immediately stored in lysis buffer (White & Densmore 1992), and genomic DNA was later extracted using Qiagen DNeasy Blood and Tissue kits. DNA concentration was determined using the Qubit dsDNA BR Assay Kit and the Qubit® Fluorometer (Life Technologies) following the manufacturers protocol.

### Microsatellite development and genotyping

We developed 12 polymorphic microsatellite loci for the variegated fairy-wren (Table S1, Supporting Information) following methods described previously in Nali *et al.* (2014). Briefly, we extracted genomic DNA from blood in lysis buffer from eight adults in our study population, enriched the mix of DNA with repetitive sequences to develop an enriched microsatellite library, and conducted an Illumina MiSeq sequencing run. From this pool of sequences, we optimized 12 loci that amplified well using polymerase chain reaction (PCR), were polymorphic, and exhibited clearly defined peaks for genotyping. We designed three multiplexed PCRs for genotyping, and each amplification reaction contained 1ul of genomic DNA of varying concentrations (1 ng/ul-40 ng/ul). PCR products were combined with the GeneScan 500 base pair LIZ internal size standard for size-sorting using a 3730 DNA Analyzer. We used Geneious version 8.0 (Kearse *et al.* 2012) to score alleles. The program automatically identifies alleles at each locus, and we manually inspected allele calls to minimize genotyping error. In total, we genotyped 287 adults and 482 nestlings from 226 nests sampled during the 2012-2016 breeding seasons.

### ddRAD sequencing

We selected a subset of the individuals genotyped for microsatellite loci (120 adults and 40 nestlings) for use in our ddRAD-seq experiment and subsequent analyses. To assess the reliability of our SNP panel for parentage and relatedness analysis, we chose representative nestlings from all the years of our study. Typically, we selected one nestling from any individual nest. In a few cases, we selected two nestlings that prior microsatellite analysis had assigned to the same mother but different fathers. For each nestling, we included the mother, the social father, and the genetic father as assigned by previous microsatellite analysis. Our pool of candidate parents included 24 mothers and 78 putative fathers, and 18 randomly selected individuals of both sexes, for a total of 120 adults.

Our ddRAD-seq protocol is adapted from Peterson *et al.* (2012) (see Supporting Information for a detailed protocol). Briefly, for each individual, 100ng - 500ng of DNA (20ul of DNA between concentrations of 5ng/ul - 25ng/ul) were digested with either SbfI and MspI, or SbfI and EcoRI (NEB), and ligated with one of 20 P1 adapters (each containing a unique inline barcode) and a P2 adapter (P2-MspI or P2-EcoRI). After digestion and ligation these samples were pooled in groups of 20 (each with a unqiue P1 adpater) and purified using 1.5X volumes of homemade MagNA made with Sera-Mag Magnetic Speed-beads (FisherSci) as described by Rohland & Reich (2012). Fragments of between 450 bp and 600 bp were selected using BluePippin (Sage Science) by the Cornell University Biotechnology Resource Center (BRC). Following size selection, index groups and Illumina sequencing adapters were added by performing 11 PCR cycles with Phusion DNA Polymerase (NEB). These reactions were cleaned up with 0.7x volumes of MagNA and pooled in equimolar ratios to create a single library for sequencing on one lane of Illumina HiSeq 2500 (100 bp single end, performed by BRC). The sequencing was performed with a ~10% PhiX spike-in to introduce diversity to the library.

By replicating 20 samples (run as a separate index group) with a wider BluePippin size selection range of 400-700 bp, we explored the inherent trade-off between the number of samples sequenced on a lane at a given coverage threshold and the number of SNPs recovered per sample. We similarly explored the effects of using a less frequent restriction enzyme digest (by substituting the 6 bp cutter EcoRI in place of the 4 bp cutter MspI) to generate a smaller total number of fragments in our size range, which in turn should increase the sequencing coverage of the loci screened.

To assess repeatability across index groups, we replicated two index groups, each comprised of 20 samples: one index group from the standard protocol, and the index group generated with the rare-cutter EcoRI enzyme (Table 1). We multiplexed the resulting 240 samples in 12 index groups (with 20 individuals each) which were pooled into a final library that was sequenced on one lane of an Illumina HiSeq 2500, producing 220,300,739 100 bp single end reads. Eight index groups included the samples used for assigning paternity and estimating relatedness, and four additional index groups were included to assess the variation in the number of loci recovered with changes in our molecular protocol (choice of enzyme and size selection window), and to assess repeatability (see Table 1). To avoid an additional source of variation these last four index groups each included the same 20 individuals.

**Table 1:** Experimental design. Index groups 9-12 were included in our sequencing run to assess how changes in our molecular protocol impacted the number of loci recovered. Therefore, we selected the same 20 individuals for these four index groups to reduce the possible sources of variation.

| Number of samples (index groups) | Enzymes | Size selection interval | Group |
| --- | --- | --- | --- |
| 160 samples (8 index groups, index 1-8) | Sbfl - MspI | 450 - 600 bp | All |
| 20 samples (Index 1) | Sbfl - MspI | 450 - 600 bp (replicate of above) | A |
| 20 samples (Index 9) | Sbfl - MspI | 450 - 600 bp (replicate of above) | B |
| 20 samples (index 10) | Sbfl - MspI | 400 - 700 bp (wide size selection) | C |
| 20 samples (index 11) | Sbfl - EcoRI | 450 - 600 bp (infrequent 3' cutter) | D |
| 20 samples (index 12) | Sbfl - EcoRI | 450 - 600 bp (replicate of above) | E |

### SNP data analysis

#### Quality filtering and demultiplexing

After the quality of the reads was assessed using FASTQC version 0.11.5 (www.bioinformatics.babraham.ac.uk/projects/fastqc), we trimmed all sequences to 97bp using *fastX_trimmer* (FASTX-Toolkit) to exclude low quality calls near the 3’ of the reads. We subsequently removed reads containing at least a single base with a Phred quality score of less than 10 (using *fastq_quality_filter).* We additionally removed sequences if more than 5% of the bases had a Phred quality score of less than 20. Using *process_radtags* module from the STACKS version 1.37 pipeline (Catchen et al. 2013), we demultiplexed the reads to obtain files with sequences that were specific to each individual.

#### De novo assembly of RAD loci

Because we do not have a sequenced genome for the variegated fairy wren or a close relative – which is likely to be the case for many non-model organisms involved in parentage studies – we assembled the sequences de novo using the STACKS pipeline (Catchen *et al.* 2013). If the genome of the species of interest (or a closely related species) is available, a reference-based assembly of RAD loci is preferred (Shafer et al. 2017). First, we used *denovo_map.pl* to assemble the reads into a catalog allowing a minimum stack depth of 5 (m parameter), up to 5 mismatches per locus within an individual (M paramenter), and 5 mismatches between loci of different individuals when building the catalog (n parameter). This combination of parameters has been shown to work well for other similarly polymorphic passerine birds (Campagna *et al.* 2015), but we expect the optimal set of parameters to vary across datasets. For a detailed exploration on how the different assembly parameters in STACKS impact the number and quality of loci recovered, see Paris *et al.* (2017). We ran the *rxstacks* module to filter loci with a log likelihood of less than −50 (lnl_lim-50) or that were confounded in at least 25% of the population (conf_lim 0.25). We then built a new catalog by rerunning *cstacks* and obtained individual genotype calls with *sstacks.*

#### SNP filtering

SNPs were exported using the *populations* module of the STACKS pipeline. All of our samples were grouped in one population and a locus was exported if it was present in 95% of the individuals in this population (r parameter) at a stack depth of at least 10 (m parameter). When a RAD locus had more than one SNP, the data were restricted to the first one *(--write_single_snp)* to avoid including SNPs in high linkage disequilibrium (LD). We required a minor allele frequency of at least 0.25 to process a nucleotide site (--*min_maf*).

We removed loci that were not in Hardy-Weinberg equilibrium using VCFtools version 0.1.14 (Danecek *et al.* 2011; ---hwe, α = 0.05). VCFtools implements an exact test which attempts to control for type I error in large datasets (Wigginton et al. 2005). VCFtools was also used to test for LD by calculating r^2^ values for every pairwise combination of the 411 SNPs in the final dataset. This analysis confirmed that LD was very low (average of 0.01; highest values: 0.43, 0.5 and 0.65). We decided to retain the entire dataset for further analyses because of the overall low LD values across all pairwise combinations. LD could be more pronounced when using enzymes that cut at higher frequencies, therefore it is advisable that LD be assessed before completing downstream parentage analyses. We obtained a variant call format (vcf) file that was converted to GENEPOP format in PGD Spider version 2.0.5.0 (Lischer & Excoffier 2012) and imported into CERVUS version 3.0.7 (Kalinowski *et al.* 2007).

#### Assessing repeatability and overlap in different sets of RAD loci

We conducted five independent de novo assemblies using STACKS, one for our larger set of 160 samples, and one for each of the four index groups included in the sequencing run to assess repeatability and understand the impact of variations in the protocol on the number of loci recovered (Table 1). Once the assemblies were completed, we filtered and exported loci independently as described above. We then queried what the overlap among these sets of loci was (e.g., between the full set of 160 samples and a replicate of 20 samples – index 9). To better understand sources of variation in our experimental protocol we also asked if the loci retained after filtering were present in the catalogs of different assemblies. To assess overlap in sets of RAD loci we first generated FASTA files with the sequence data from both the filtered loci from each assembly and from the entire catalog. We then used BLAST version 2.3.0 (Altschul *et al.* 1990) with an E-value of 1e-10 to find matches among these FASTA files and calculate the proportion of RAD tags shared by the different sets of sequences (Tables 2 and 3).

**Table 2.** Overlap in the RAD loci that were obtained while varying different steps of the protocol (size selection and restriction enzymes). The diagonal indicates the total number of loci recovered for each treatment. Values above the diagonal represent the percent overlapping loci between groups (relative to the group with the smallest number of loci), while values below the diagonal list the number of loci that were overlapping between groups.

|  | **All** | **A** | **B** | **C** | **D** | **E** |
| --- | --- | --- | --- | --- | --- | --- |
| **All** | 411 | 85.4 | 78.1 | 79.1 | 1.2 | 1.0 |
| **A** | 351 | 797 | 85.1 | 75.8 | 0.8 | 0.8 |
| **B** | 321 | 549 | 645 | 83.4 | 2.0 | 2.3 |
| **C** | 325 | 604 | 538 | 1440 | 4.2 | 3.9 |
| **D** | 5 | 15 | 12 | 25 | 596 | 68.4 |
| **E** | 4 | 13 | 12 | 20 | 353 | 516 |

All: 160 samples; Sbf1/Msp1; 450-600 bp. A: 20 samples; Sbf1/Msp1; 450-600 bp (Index 1). B: 20 samples; Sbf1/Msp1; 450-600 bp (Index 9). C: 20 samples; Sbf1/Msp1; 400-700 bp (Index 10). D: 20 samples; Sbf1/EcoRI; 450-600 bp (Index 11). E: 20 samples; Sbf1/EcoRI; 450-600 bp (Index 12).

**Table 3.**
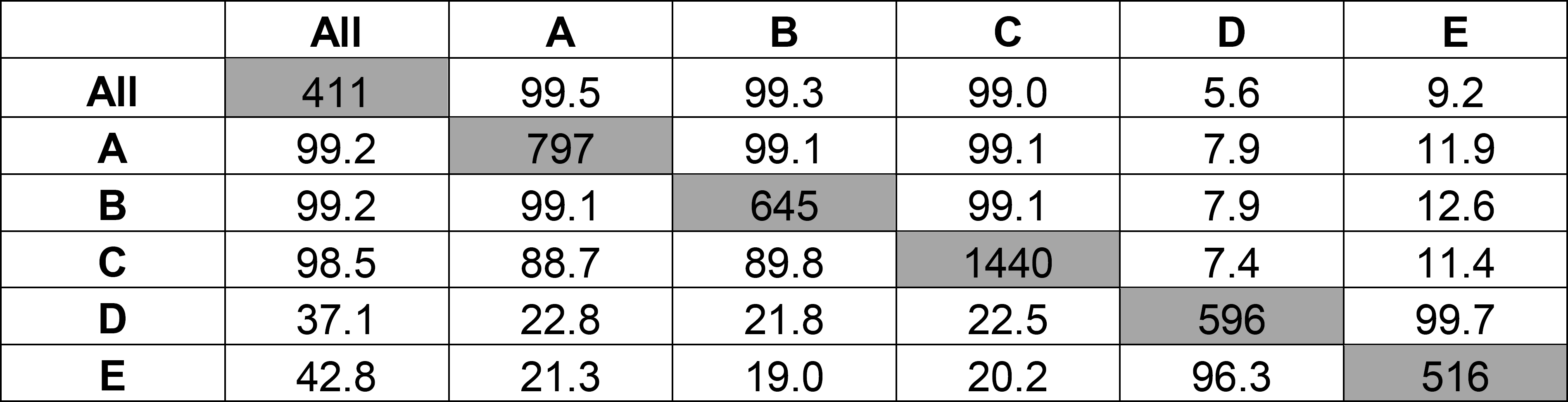
Overlap in filtered RAD loci and those present in the catalogs from the different assemblies. The filtered loci from the groups in the different rows were aligned against databases generated from the catalogs of the groups in the columns. The numbers in the table are the percent of filtered loci from each group that match loci that are in the catalog of the target database. The diagonal contains the total number of loci recovered for each group and the total number of loci in each catalog are 49662 (All), 27085 (A), 25521 (B), 29554 (C), 11823 (D), 16724 (E).

|  | **All** | **A** | **B** | **C** | **D** | **E** |
| --- | --- | --- | --- | --- | --- | --- |
| **All** | 411 | 99.5 | 99.3 | 99.0 | 5.6 | 9.2 |
| **A** | 99.2 | 797 | 99.1 | 99.1 | 7.9 | 11.9 |
| **B** | 99.2 | 99.1 | 645 | 99.1 | 7.9 | 12.6 |
| **C** | 98.5 | 88.7 | 89.8 | 1440 | 7.4 | 11.4 |
| **D** | 37.1 | 22.8 | 21.8 | 22.5 | 596 | 99.7 |
| **E** | 42.8 | 21.3 | 19.0 | 20.2 | 96.3 | 516 |

### Parentage analysis

We used CERVUS version 3.0.7 (Kalinowski *et al.* 2007) to assign paternity for all nestlings using our microsatellite and SNP datasets separately. CERVUS uses a two-step, likelihood-based approach to assign parentage. First, CERVUS compares each offspring’s genotype to that of a candidate parent and a random individual in the population to calculate a likelihood ratio. This relationship is presented as an LOD score, which is simply the natural logarithm of the calculated likelihood ratio. Positive LOD scores indicate that a candidate parent is much more likely to be the true parent, whereas negative LOD scores indicate that the candidate parent is highly unlikely to be a true parent. Second, CERVUS conducts a simulation of parentage analysis based on population allele frequencies and the proportion of potential parents included in the analysis. The simulation accounts for the possibility of unsampled parents, missing data, and genotyping errors. Considering these parameters, the simulation calculates critical LOD scores by comparing the LOD distributions of the most likely parent and all other candidate parents. The critical LOD score is used to determine the confidence (95% or 80%) of each parentage assignment.

CERVUS allows for different types of parentage analysis, including parent-pair (sexes known or unknown), maternity (known father, but not mother), and paternity (known mother, but not father). Variegated fairy-wrens at Lake Samsonvale are relatively easy to observe, and we were able to assign known mothers behaviorally. We subsequently confirmed this with microsatellite analyses: females that built and attended a nest throughout incubation were always the mothers of the nestlings in that nest. In many systems, a comparable level of demographic knowledge may not be available, so a marker set must be powerful enough to assign parentage with minimal social information. To investigate the broader utility of our ddRAD-seq method, we conducted analyses that relied on the inclusion of the known mother, in addition to analyses independent of the known mother, which were based only on the father-offspring relationship. We simulated paternity assignments for 10,000 offspring to determine critical LOD scores, using slightly different input parameters for each panel (microsatellites and ddRAD sequencing derived SNPs). Simulations for both used the following parameters: 78 candidate males, 95% of candidate males sampled, estimated error rate of 0.01 for mistyped loci and likelihood scores. The proportion of loci typed across all individuals was different for both panels: 0.997 for the microsatellite simulation, and 0.961 for the SNP simulation.

For both paternity analyses, we used the trio LOD score and the father-offspring LOD score from CERVUS to make assignments. The trio LOD score was calculated by comparing the genotypes of the candidate male and offspring, relative to that of the known mother. The father-offspring LOD score only accounts for the relationship of the candidate male and the offspring, independent of the known mother. CERVUS ranked candidate males by LOD scores in each category, and the highest-ranking males were assigned as fathers. These rankings should be in agreement, but ambiguous assignments (different top-ranking males assigned in each category) may occur when multiple candidate male genotypes closely match an offspring’s genotype.

We assessed each CERVUS assignment to determine whether it was plausible, and whether the assigned male was the social father or an extra-pair sire. Our criteria for accepting assignments differed slightly for microsatellites and SNPs. For microsatellites, we automatically accepted the CERVUS assignment if the highest-ranking male was in agreement for both the trio LOD and the father-offspring LOD, and if the number of mismatches between the assigned male and the offspring was < 1 (8% of 12 loci). For SNPs, we also accepted the assignment if the highest-ranking males by LOD score type were in agreement, and an allowable number of mismatches were not exceeded. However, for SNPs, our allowable number of mismatches was based on the observed maximum number of mismatches between a known mother and her known offspring (max. = 7, mean = 3.4, 2% of 411 loci). For both panels, we accepted the social father as the genetic sire if he met these respective criteria. If the social father mismatched the offspring at higher numbers, or had negative LOD scores, the offspring was considered sired by an extra-pair father. We accepted assignments of extra-pair fathers using the same criteria outlined above. We did not observe cases in which an offspring could not be assigned to either its social father, or an extra-pair sire.

### Relatedness analysis

We used the package, ‘related’ (Pew *et al.* 2015), in R version 3.2.5 (R Core Team 2016) to estimate pairwise relatedness (r) between all pairs of individuals in this study. This package accounts for genotyping errors, missing data, and can estimate relatedness using any of seven different estimators (4 non-likelihood-based, and 3 likelihood-based). ‘Related’ includes the function, *compareestimators,* which tests the performance of different estimators on simulated data that share the same characteristics as the real data. The program uses an allele frequency file to generate simulated pairs of individuals of known relatedness, and automatically estimates relatedness using four of the most commonly used estimators (all non-likelihood-based). The function calculates a correlation coefficient between observed and expected values, to evaluate which estimator performs best with the data set. Using *compareestimators* to generate 200 simulated pairs of individuals for each degree of relatedness (i.e., half-sib, full-sib, parent-offspring, unrelated), we determined that the Wang (2002) estimator performed best for both our microsatellite and SNP datasets. However, SNP datasets generated by ddRAD-seq are sometimes prone to genotyping error through allelic dropout. Attard *et al.* (2018) found that this can cause relatedness estimators to produce values that are very precise, but slightly downward-biased, especially for large datasets. This should be considered when selecting an appropriate relatedness estimator for standalone SNP data. For consistency across the comparisons of our microsatellite and SNP datasets, we obtained point estimates of relatedness using the Wang (2002) estimator, and evaluated all parent-offspring relationships that were previously determined in our parentage analysis.

‘Related’ also evaluates how well different marker sets resolve degrees of relatedness, given simulated genotypes based on allele frequency files. For both panels, we used the *familysim* function to generate 200 pairs of individuals for each degree of relatedness. We then used the *coancestry* function to analyze all pairwise relatedness values with the Wang (2002) estimator. We created density plots representing histograms of the relatedness values. These plots show the overlap in relatedness values for degree of relatedness, and we used them to infer how well each panel performed at discerning different relationships.

### Comparison with other avian ddRAD datasets

We applied the same molecular protocol and bioinformatics pipeline described above to other avian species, with the objective of assigning paternity and estimating relatedness. These datasets ranged between 6 and 480 samples, and we calculated the number of loci recovered for comparisons with the current data and to assess the utility of our method with larger sample sizes. We also genotyped an additional 213 variegated fairy-wren individuals and re-analyzed the data in combination with the 160 samples included in this study.

## Results

### SNP development and analysis

After trimming, filtering and demultiplexing the data, we retained a total of 109,524,874 reads across all index groups, with an average of approximately 9,000,000 reads per index group. Two individuals failed (i.e., had less than 66,000 reads each; one individual from each of two index groups) and were excluded from further analysis. The number of reads per sample for the remaining individuals ranged from 213,544 to 810,966 (mean = 459,726 ± 118,771 std. dev.).

Further analysis using the population program from STACKS identified loci with at least 10X coverage, present in 95% of the individuals, and a minimum allele frequency greater than 25%. We further retained only those loci that were in Hardy-Weinberg equilibrium. When performing analyses on all 160 individuals from the primary comparison runs (Table 1), we identified 411 loci that fulfilled these criteria and were used for downstream analyses.

We varied two aspects of the ddRAD-seq protocol to assess the number of loci recovered and the reproducibility of the method. As expected, both a reduction in the range of fragment sizes selected during the construction of the library (compare groups A (150 bp size selection) and C (300 bp size selection)), or the use of the less frequent cutter, EcoRI (groups D and E), resulted in fewer loci recovered (Table 2). Those from the EcoRI digest were mostly non-overlapping with those from the MspI index groups. More importantly, between the replicated index groups in our standard protocol (A and B) we recovered similar numbers of loci (797 and 645, respectively) with 549 (85.1%) loci found in both datasets under our stringent filtering criteria. Moreover, when we searched for the loci recovered in A in the catalog of B, and vice versa, the overlap was above 99% (Table 3). We also found that 208 of the 248 non-overlapping loci recovered in A were not recovered in B because they did not pass the stringent missing data (r = 0.95) and minimum depth of coverage (m = 10) filters (Table S2, Supporting Information). This suggests that very similar sets of loci were recovered in these independent assemblies, but that there was variation in the loci that passed our filtering criteria. Although changing our filtering parameters led to a slight increase in the proportion of overlap in the loci recovered between A and B, the total number of overlapping loci increased substantially. When we accepted 20% missing data instead of 5%, the proportion of overlap was 86%, with 857 loci in total. When we changed the minor allele frequency filter from 0.25 to 0.05 we obtained 88% overlap and a total of 1227 loci (Table S3, Supporting Information).

### Paternity assignments

Both panels produced highly concordant results when assigning paternity, but the SNP panel showed substantially higher power overall. Generally, the microsatellite loci were more polymorphic, resulting in greater mean polymorphic information content (PIC) for any given locus. Despite this, the SNP panel performed better because of the large number of loci obtained through RAD sequencing. This greatly improved the non-exclusion probabilities across different parentage assignment contexts (Table 4), and reduced uncertainty in our assignments. Given the known mother, the microsatellite and SNP panels assigned the same fathers to all 40 offspring with 95% confidence. When paternity assignments were made without the known mother (no known candidate parents), both panels again assigned fathers for all 40 offspring with 95% confidence. However, 5 of these assignments were not in agreement between the two panels. For these 5 cases, two candidate males had very similar LOD scores under the microsatellite panel, and the assigned males did not match the males that were assigned given the known mother. These (and all other) cases were resolved unambiguously when the SNP panel was used, and the paternity assignments with and without the known mother were in complete agreement for all offspring (Fig. 1).

**Table 4.** Marker characteristics. He: Expected heterozygosity; Ho: Observed heterozygosity; PIC: Polymorphic information content.

| Marker Panel | Number of loci | Mean proportion loci typed | Mean alleles per locus | Mean $H_e$ | Mean $H_o$ | Mean PIC | Nonexclusion probability (first parent) | Nonexclusion probability (second parent) | Nonexclusion probability (parent pair) |
| --- | --- | --- | --- | --- | --- | --- | --- | --- | --- |
| Microsatellites | 12 | 0.99 | 14.17 | 0.77 | 0.76 | 0.74 | $1.9 \times 10^{-4}$ | $1.9 \times 10^{-6}$ | $1.5 \times 10^{-10}$ |
| SNPs | 411 | 0.98 | 2.00* | 0.45 | 0.45 | 0.35 | $5.2 \times 10^{-20}$ | $6.9 \times 10^{-35}$ | $1.0 \times 10^{-55}$ |

\*Only biallelic SNPs were retained. If a locus had 3 alleles across the population, it was filtered from the dataset.

**Figure 1.**
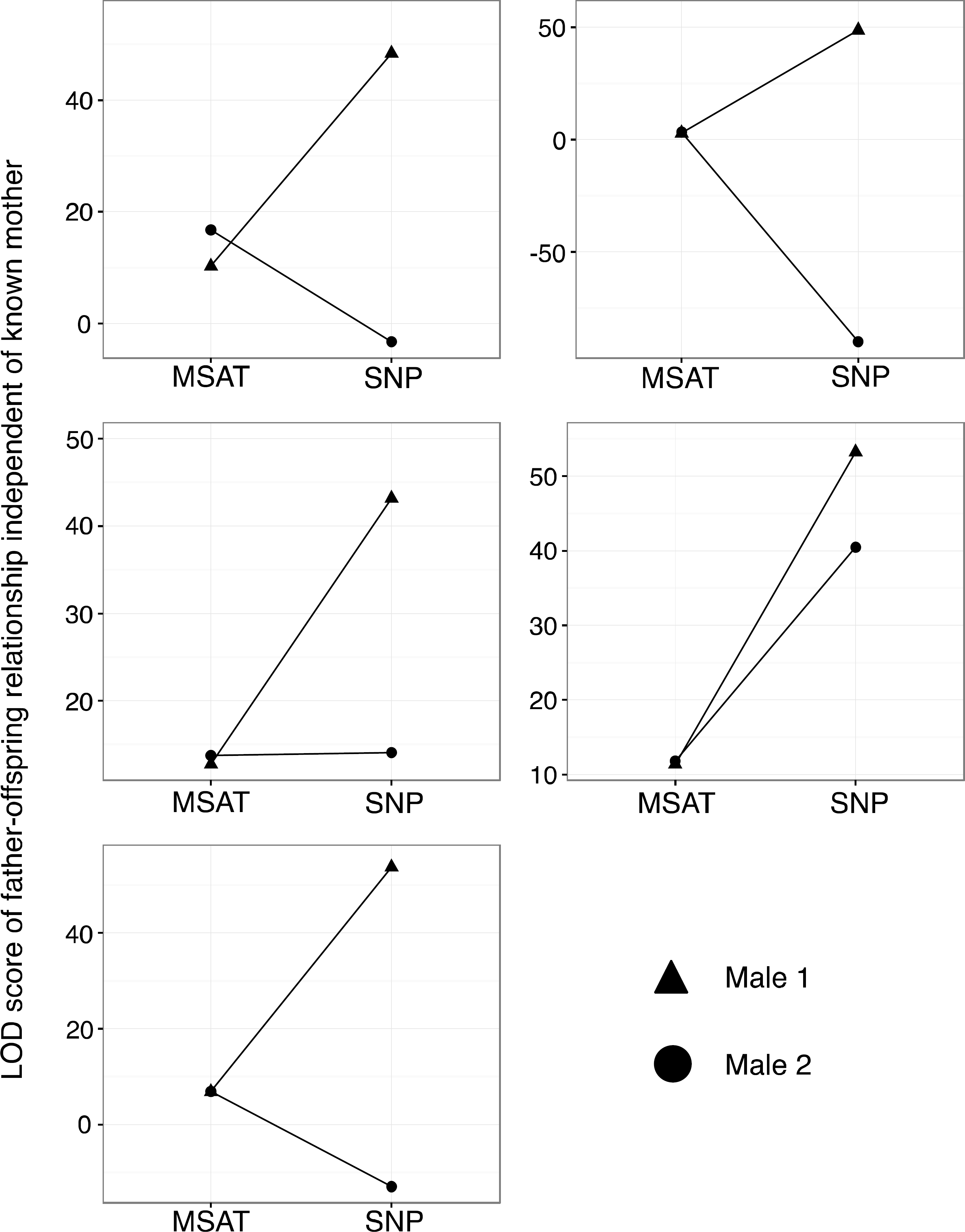
Resolved paternity assignments for 5 nestlings with ambiguous assignments under the microsatellite panel, but not with the SNP panel. Each panel in the graph represents an individual offspring, and the two top-ranked males are depicted as a triangle and a circle, respectively. Lines connecting like shapes show the change in LOD score for each male, using each marker type (microsatellites versus SNPs). Note that the y-axis scale varies among panels in the graph.

Overall, both panels assigned 23 out of 40 nestlings (57.5%) to males that were not their social father. Due to the nature of our non-random sampling of individuals for this experiment, and the overall smaller sample size, this value is slightly lower than the overall rate of 67.6% extra-pair young observed for all years of the study (unpublished data).

As a measure of certainty for our assignments, we calculated the difference between LOD scores for the two top-ranked males assigned to each nestling, under each panel (Fig. 2). Typically, this difference was 8 – 10x higher for the SNP panel (n=40, mean = 165.0) than for the microsatellite panel (n=40, mean 19.1), a reflection of the much higher discriminatory power of the SNP dataset. For the SNP panel, many of the second-ranked males had a strongly negative LOD score, making them extremely unlikely to be the true father. This was less often true for the microsatellite panel, as the second-ranked males often had positive, or just slightly negative, LOD scores. Overall this result illustrates the increased discrimination power achieved by the SNP panel compared to the microsatellites, which allowed us to assign paternity in cases in which the microsatellite assignments remained ambiguous or (albeit rarely) misleading.

**Figure 2.**
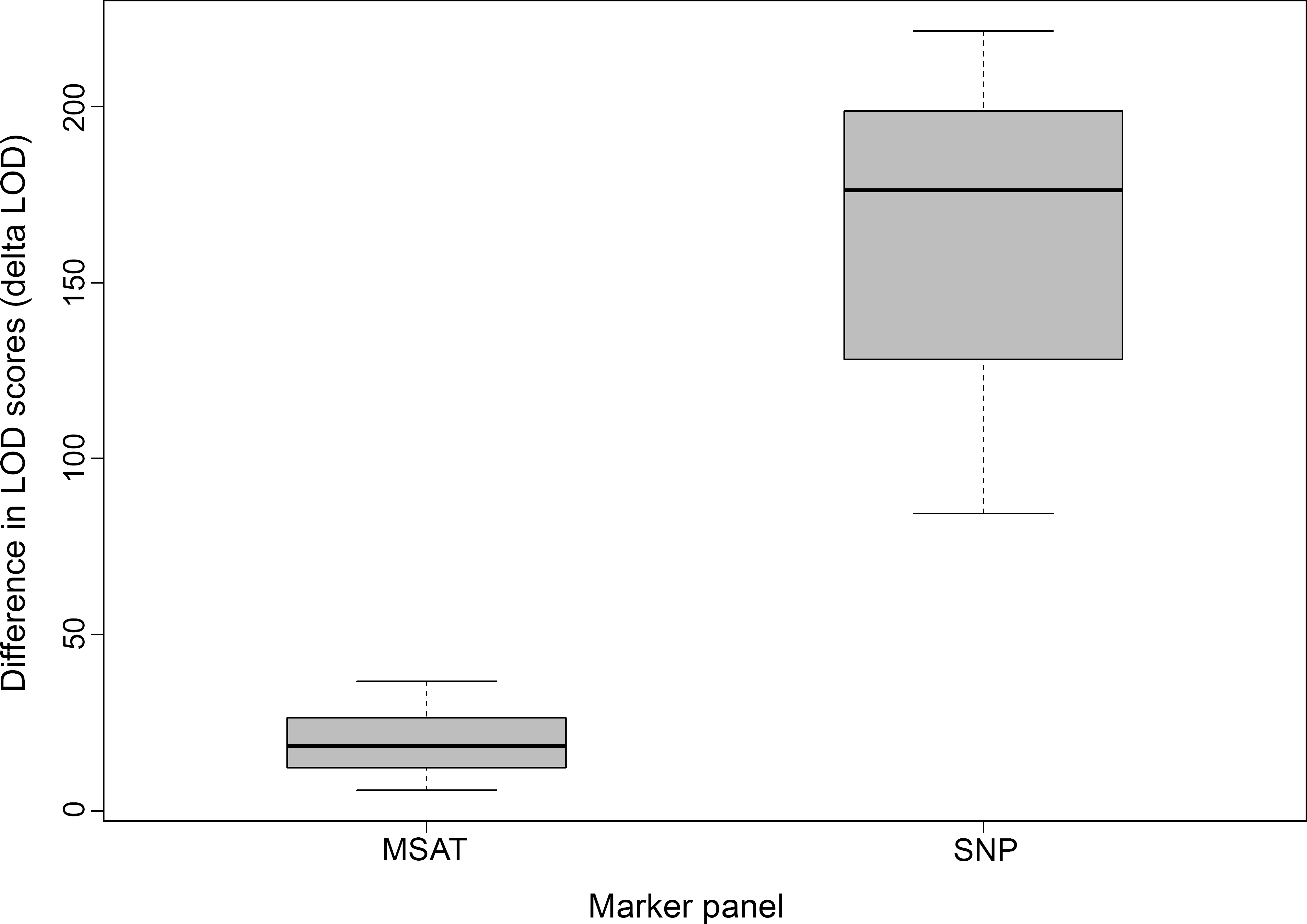
Difference in CERVUS LOD scores (delta LOD) between the most likely father of a nestling and the second possible father in the population, for both marker panels.

### Relatedness analysis

The SNP panel produced simulated data that closely matched the observed allele and genotype frequencies (Pearson’s correlation coefficient = 0.975). The microsatellite panel also matched well, but was not as reliable as the SNP panel (Pearson’s correlation coefficient = 0.877). This resulted in better estimates of pairwise relatedness for parent-offspring using the SNP panel (Fig. 3). Overall, the SNP panel produced better simulated estimates for each degree of relatedness (Fig. 4), greatly reducing the variance around expected relatedness values (unrelated = 0, half-sib = 0.25, full-sib = 0.5, and parent-offspring = 0.5). This bolsters the confidence with which actual relationships can be discerned when calculating pairwise relatedness of a population for which there is little prior knowledge of social relationships.

**Figure 3.**
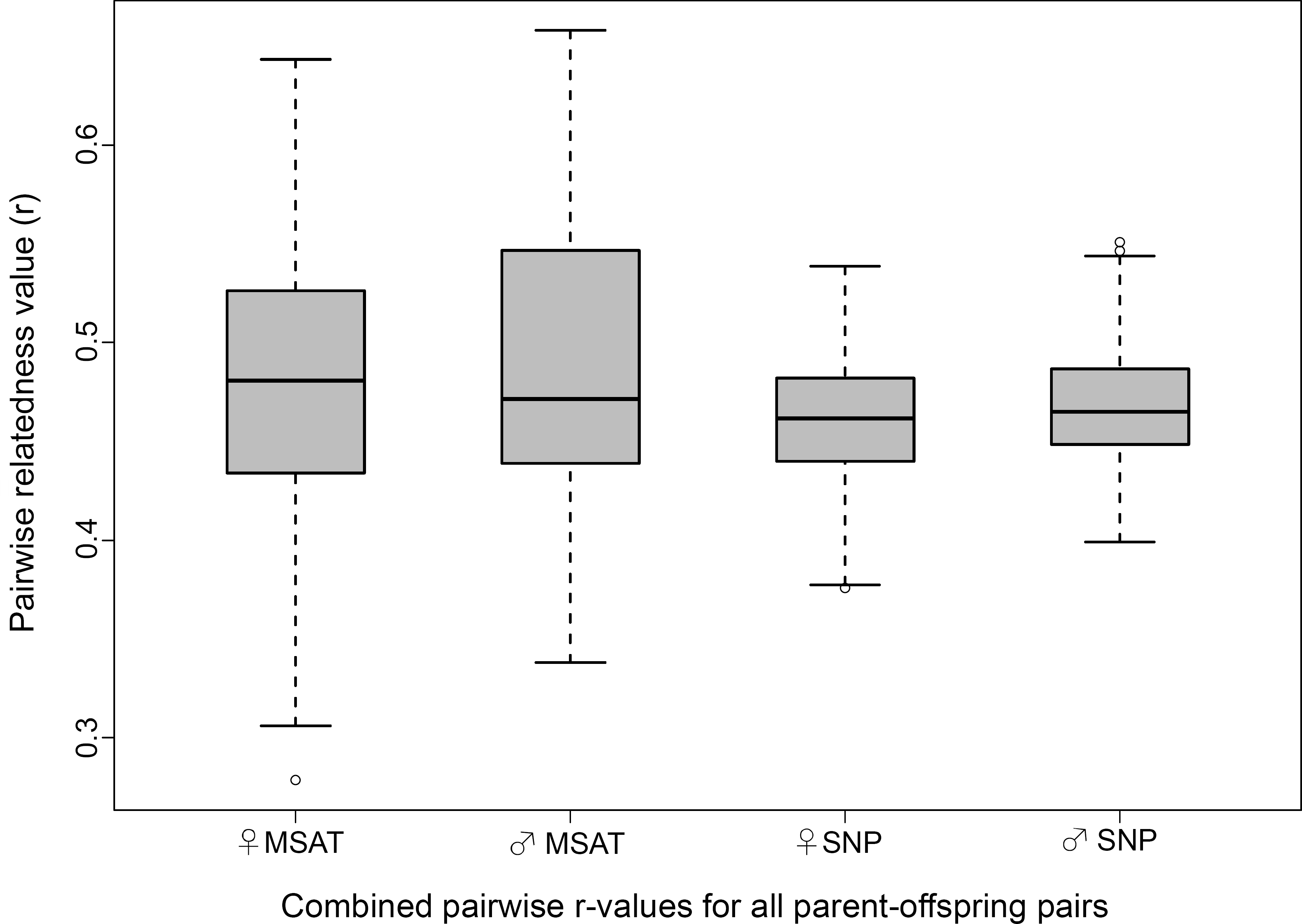
Box plot of pairwise relatedness values for all parent-offspring (40 mother-offspring and 40 father-offspring) relationships, using population allele frequencies from each marker panel.

## Discussion

Several recent studies have rigorously investigated the use of SNPs in population genetic studies for several non-model organisms (Morin *et al.* 2004; Slate *et al.* 2010; Garvin *et al.* 2010; Heylar *et al.* 2011; Seeb *et al.* 2011), with growing support for the use of SNPs in studies of parentage (Anderson & Garza 2006; e.g. Hauser *et al.* 2011; e.g. Kaiser *et al.* 2017; e.g. Kess *et al.* 2016) and relatedness (e.g. Glaubitz *et al.* 2003; Wang 2007). SNPs have proven to perform as well, if not better than microsatellites in these types of studies. To our knowledge, this is the first study to describe a ddRAD-seq method for use in parentage and relatedness analyses of wild populations, and to test it in a diverse set of avian taxa. Our study is also the first to compare the efficiency of microsatellites versus SNPs for determining genetic relationships in a bird species that is both socially complex and highly promiscuous. We show that SNPs developed from our modified ddRAD-seq method are substantially more powerful than a moderate number of species-specific microsatellite loci at assigning paternity and estimating relatedness among individuals. Our method is highly attractive as an alternative to traditional microsatellite genotyping, especially for systems where no microsatellites have been developed. This is largely due to the combination of its cost and researcher time efficiency, the ease of this non-species-specific method that combines the SNP discovery and screening steps, and the large number of SNPs reliably recovered.

The total approximate materials cost for our ddRAD-seq analyses, including DNA extraction, normalization of the DNA concentrations, library preparation, sequencing and computational time was US $3,270.00 for 240 samples, or approximately $13.6 per sample. The initial investment in oligonucleotides (i.e., primers and adaptors; see the Supplementary ddRAD-seq Protocol in the Supporting Information) was US $2000 and is sufficient for the analysis of thousands of samples, making the per sample cost negligible. The use of a homemade MagNA in place of commercial SPRI beads provides significant savings. This cost is similar to that for genotyping 240 individuals at 12 microsatellite loci (in 3 multiplexed PCR mixes), in a situation where the labelled primers have already been designed, purchased, and tested. However, a substantial additional benefit of this ddRAD-seq method is that it does not require any locus discovery or development before starting. The time required for library preparation, once DNA has been extracted, is modest, and once the sequence data have been obtained, SNP calling for the entire dataset can be performed in less than a day through a largely automated bioinformatics pipeline (see Supporting Information). Unlike manually scoring peaks in traditional microsatellite genotyping analyses, the identification of SNPs is less subjective and takes far fewer hours of hands-on analysis (as most is performed computationally). The tools for analyzing these ddRAD data are freely available and widely used (e.g., STACKS, VCFtools). Nevertheless, we note that assembling RAD loci can still be challenging, and the choice of bioinformatics pipelines and specific combinations of assembly parameters can influence the quality and quantity of loci recovered (for detailed discussions on these issues see Eaton 2014, Mastretta-Yanes *et al.* 2015, Shafer *et al.* 2017 and Paris *et al.* 2017).

For this study, our conditions and protocol allowed us to recover 411 high quality SNP loci for 160 individual samples (although 240 samples were multiplexed together on one lane of sequencing). However, we show that through simple variations in the size selection window or the specificity of the restriction enzyme, more or fewer loci can be obtained. For some applications, it could be advantageous to multiplex a greater number of individuals and achieve similar coverage by aiming to recover fewer loci (e.g., using EcoRI rather than MspI). Alternatively, for applications where more loci are required, the size selection window could be widened and concordantly the number of individuals would have to be lowered. It is also possible to vary the number of loci retained by applying different bioinformatics filters. With strict filtering parameters (5% missing data, minor allele frequency of 0.25 and minimum depth of coverage of 10x) we recovered 411 loci, which contained sufficient information to accurately assign paternity and estimate relatedness among the individuals in our study. However, with only slight modification of these parameters it is possible to greatly increase the number of loci recovered. A total of 506 loci were retained when we allowed a minimum coverage of 5x, 742 with up to 20% missing data, and 910 with a minor allele frequency of 0.05 (we varied one filter at a time). With respect to the entire catalog from 160 individual samples (49662 loci from the ALL group), when we applied one filter at a time, we found that approximately 47% of loci remained when a minimum coverage of 10x was required, while only 5% remained when we allowed up to 5% missing data (Table S4, Supporting Information). This suggests that the greatest loss of loci across all individuals can be attributed to missing data, and to a lesser extent, depth of coverage. These missing loci may be the result of DNA quality, conservation of restriction sites, size selection, or per sample coverage during sequencing. Despite these losses under our stringent filtering criteria, we still recovered more than a sufficient number of informative SNPs to perform highly robust parentage and relatedness analyses.

The number of SNPs needed to perform robust parentage and relatedness analyses depends on characteristics of the study population. Populations with reduced genetic diversity will likely require a greater number of loci than those that are more genetically diverse (Saunders *et al.* 2007; Strucken *et al.* 2016; Tortereau *et al.* 2017). Obtaining more loci from the outset would aid in overcoming any issues relating to population genetic diversity. Additionally, when studying species with complex social systems, including for example both variable levels of genetic relatedness among individuals and high rates of extra-pair fertilizations, it is imperative to obtain a sufficient number of markers to discern genetic relationships robustly (Hughes 1998; Ross 2001; Weinman *et al.* 2015). Our case study, using the variegated fairy-wren, shows that our modified ddRAD-seq method recovers more than enough SNP loci to confidently discern relationships in a species with a complex social system. Most parentage and relatedness analysis programs are well equipped to handle large numbers of loci, so a greater number of loci would not hinder analyses. Once an appropriate number of SNPs are identified for performing robust analyses, conditions can be varied to maximize the number of individuals to be genotyped. For our purposes, we conducted parentage and relatedness analyses in CERVUS and the R package ‘related,’ respectively, to reliably compare the performance of our microsatellite and SNP panels. Several other pedigree reconstruction programs are readily available (e.g. COLONY, MasterBayes, and Sequoia) and researchers can easily input SNP data into their preferred program (for detailed comparisons of some of these programs see Karaket *et al.* 2012 and Weinman *et al.* 2015). The R package, ‘Sequoia’ (Huisman 2017), is specifically tailored for SNP data, and can reconstruct multi-generational pedigrees with as few as 100 SNPs and many non-genotyped individuals. Given these considerations, ‘Sequoia’ may be particularly useful for studies with limited social information or incomplete population sampling.

For both paternity and relatedness analyses, our SNP panel far outperformed our microsatellite panel by providing much more power and improving the overall confidence for assignments. Variegated fairy-wrens are relatively easy to observe, and every nest found can be assigned to a known mother by watching the female that builds the nest and/or incubates the eggs. This level of knowledge may not be the norm for most study systems, so we also investigated the CERVUS output for male-offspring relationships, independent of known mothers. In doing so, the reliability of the SNP panel became even more evident. In CERVUS, the higher the LOD score, the more likely that a given male is the true father. Using SNPs, CERVUS typically output only a single male with a positive LOD score, and the difference in LOD scores between the top-two ranked males was dramatically different for SNP assignments (Fig. 2). When social information about the known mother was excluded from the paternity analysis, the microsatellite panel sometimes produced assignments that were ambiguous (two males had similar LOD scores), and occasionally the wrong male was assigned paternity of the offspring. Under the SNP panel, ambiguous assignments were nonexistent, and these cases were clearly resolved (Fig. 1).

It is sometimes difficult to obtain appropriate demographic data to use in a formal parentage analysis, and for many studies, this level of detail may not be necessary. Population allele frequencies can be used to estimate pairwise relatedness for individuals, and to reconstruct pedigrees using maximum likelihood-based methods. Variance in estimates of pairwise relatedness (r) for known parent-offspring pairs was dramatically reduced when using SNPs (Fig. 3). For our simulations, SNPs greatly improved the differentiation between distributions for individuals of known degrees of relatedness (Fig. 4). This is particularly important for systems with minimal demographic and observational data, where these distributions can be used to determine familial relationships between individuals, in conjunction with actual estimated r-values.

**Figure 4.**
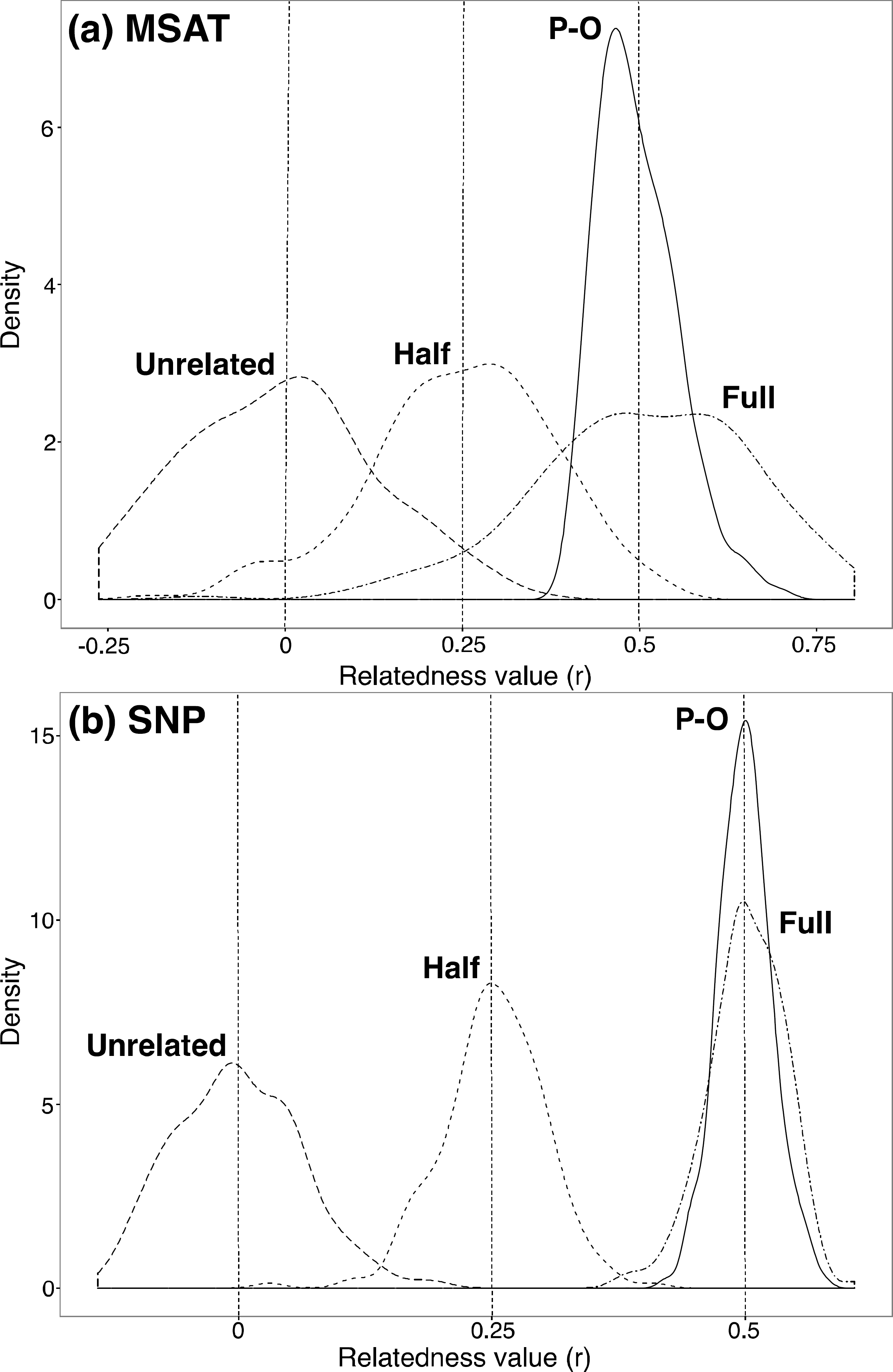
Density plots of relatedness values for simulated pairs of known relatedness (unrelated, half-sibling, full-sibling, and parent-offspring) using population allele frequencies from each marker panel (a. MSAT; b. SNP). Overlap in distributions indicates the overlap between relatedness value estimators for pairs of individuals of different relationships. The spread of each distribution indicates the reliability of observed relatedness values based on their deviation from expected relatedness values (Unrelated = 0, Half-sib = 0.25, Full-sib = 0.5, and Parent-offspring (P-O) = 0.5, denoted by vertical dashed lines).

This protocol was designed to be universally applicable across bird species, and we have successfully applied it in a range of other avian study systems (Table 5). While different numbers of individuals were used in each study, and therefore different numbers of loci were recovered, in all cases paternity was confidently assigned to nestlings using CERVUS (unpublished results). We note that when the number of individuals genotyped was larger (we have so far tested up to 480 individuals), the number of loci recovered after filtering was smaller (as low as 135 SNPs with stringent filtering parameters). Our controls designed to assess repeatability (e.g., compare group A and B or D and E in Tables 2 and 3) suggest that there is a high degree of overlap among replicates, but also variation in which loci pass our filters. In cases in which a larger number of loci are needed to accurately assess paternity or estimate relatedness, some of the filtering parameters may be relaxed. However, further testing would be required to assess the informativeness of loci obtained under less stringent filtering criteria. For example, for a set of 373 Variegated fairy-wren individuals (Table 5) the number of loci retained increased from 157 to 410 when 20% missing data was allowed instead of 5%. Accordingly, our protocol is suitable for long-term studies in which samples are accumulated across several years, especially if not all individuals need to be compared simultaneously (e.g. non-overlapping generations or individuals from different years). It is also likely that this protocol can be applied successfully for studies (short or long-term) where thousands of individuals need to be compared at one time. This remains to be shown, though, and in such cases it may be more appropriate to use techniques based on microarrays (see Fernández *et al.* 2013, Liu *et al.* 2016, and Tortereau *et al.* 2017).

**Table 5:** Summary information for ddRAD-seq studies performed to investigate parentage in other bird species. Only SNPs with up to 5% missing data, with a minimum coverage of 10x and a minor allele frequency of 0.25 were retained. HWE: Hardy-Weinberg equilibrium.

| Species | Scientific name | Number of individuals | Number of loci | Number of loci in HWE |
| --- | --- | --- | --- | --- |
| Variegated fairy-wren* | *Malurus lamberti* | 373 | 234 | 157 |
| Hispaniolan woodpecker** | *Melanerpes striatus* | 288 | 179 | 135 |
| Northern Red-billed hornbill | *Tockus erythrorhynchus* | 40 | 475 | 414 |
| Von der Decken's hornbill | *Tockus deckeni* | 112 | 490 | 410 |
| Sapayoa | *Sapayoa aenigma* | 6 | 672 | 671 |
| Red-backed fairy-wren*** | *Malurus melanocephalus* | 480 | 329 | 167 |

\* Two independent ddRAD-seq experiments were performed – one with 160 samples (those from the current study) and a second with 213 samples. After filtering and demultiplexing, the data from 373 samples were combined for denovo assembly and SNP identification.
\*\* Samples were run in two experiments and combined, one with 240 samples and the other with 48.
\*\*\* Two independent ddRAD-seq experiments were run on each set of 240 samples.

Applying this general protocol to many non-avian taxa will simply require ensuring that specific restriction enzymes and fragment size windows are chosen appropriately. The size of the genome and the number of individuals multiplexed will have to be taken into consideration to achieve the desired coverage.

In summary, our ddRAD-seq method provides a cost effective and robust way to identify SNPs for use in studies utilizing parentage and relatedness analyses. Our experiment shows that a majority of the same SNPs can be obtained across groups, using the same size selection windows and restriction enzymes. Future individuals can be genotyped and incorporated to the analysis by re-running the STACKS pipeline. Using a bird exhibiting great social complexity, and high promiscuity, we have shown that SNPs identified by ddRAD-seq are more effective at assigning paternity and estimating relatedness than highly polymorphic, species-specific microsatellite loci.

## Acknowledgements

We thank D. Baldassarre, K. Gielow, J. Welklin, and field technicians at Lake Samsonvale who assisted with field efforts for this study. We are grateful to L. Stenzler and S. Bogdanowicz for help with microsatellite discovery, development, and genotyping. D. Baldassarre, E. Greig, & A. Dalziel provided valuable discussion on the study design. We thank J. LaPergola, J. Welklin, B. Van Doren, and M. Kinnaird for providing the unpublished data shown in Table 5. We are grateful to Y. Bourgeois and two anonymous reviewers for valuable comments on previous versions of this manuscript. This research was supported by the US National Science Foundation (IOS-1353681 and DEB-1721662).

## Data Accessibility

RAD loci from de novo assembly in STACKS (Groups All and A-E), and Microsatellite and SNP genotypes: Dryad Digital Repository: doi:10.5061/dryad.c76pf34.

## Author Contributions

D.J.T designed the study, collected field data, performed microsatellite development and analysis, conducted parentage and relatedness analyses, and drafted the manuscript with help from all co-authors. B.G.B and L.C. designed the study, and performed SNP discovery and analysis. M.S.W and I.J.L. helped design the study, and secured funding.

## References

Altschul SF, Gish W, Miller W, Myers EW, Lipman DJ (1990) Basic local alignment search tool. Journal of Molecular Biology, 215, 403–410.

Anderson EC, Garza JC (2006) The power of single-nucleotide polymorphisms for large-scale parentage inference. Genetics, 172, 2567–2582.

Andrews KR, Good JM, Miller MR, Luikart G, Hohenlohe PA (2016) Harnessing the power of RADseq for ecological and evolutionary genomics. Nature Reviews Genetics, 17, 81–92.

Attard, RMC, Beheregaray, LB, Möller, LM (2018) Genotyping-by-sequencing for estimating relatedness in nonmodel organisms: Avoiding the trap of precise bias. Molecular Ecology Resources, 00, 1–10.

Avise JC, Jones AG, Walker D, Dewoody JA (2002) Genetic mating systems and reproductive natural histories of fishes: lessons for ecology and evolution. Annual Review of Genetics, 36, 19–45.

Baird NA, Etter PD, Atwood TS et al. (2008) Rapid SNP discovery and genetic mapping using sequenced RAD markers. PLoS ONE, 3, e3376.

Ball AD, Stapley J, Dawson DA et al. (2010) A comparison of SNPs and microsatellites as linkage mapping markers: lessons from the zebra finch (Taeniopygia guttata). BMC Genomics, 11, 218.

Blouin MS (2003) DNA-based methods for pedigree reconstruction and kinship analysis in natural populations. Trends in Ecology & Evolution, 18, 503–511.

Brumfield RT, Beerli P, Nickerson DA, Edwards SV (2003) The utility of single nucleotide polymorphisms in inferences of population history. Trends in Ecology & Evolution, 18, 249–256.

Campagna L, Gronau I, Silveira LF, Siepel A, Lovette IJ (2015) Distinguishing noise from signal in patterns of genomic divergence in a highly polymorphic avian radiation. Molecular Ecology, 24, 4238–4251.

Catchen J, Hohenlohe PA, Bassham S, Amores A, Cresko WA (2013) Stacks: an analysis tool set for population genomics. Molecular Ecology, 22, 3124–3140.

Coates BS, Sumerford DV, Miller NJ et al. (2009) Comparative Performance of Single Nucleotide Polymorphism and Microsatellite Markers for Population Genetic Analysis. Journal of Heredity, 100, 556–564.

Cramer ERA, Hall ML, De Kort SR, Lovette IJ, Vehrencamp SL (2011) Infrequent Extra Pair Paternity in the Banded Wren, a Synchronously Breeding Tropical Passerine. The Condor, 113, 637–645.

Danecek P, Auton A, Abecasis G et al. (2011) The variant call format and VCFtools. Bioinformatics, 27, 2156–2158.

Davey JW, Blaxter ML (2010) RADSeq: next-generation population genetics. Briefings in Functional Genomics, 9, 416–423.

Davey JW, Hohenlohe PA, Etter PD et al. (2011) Genome-wide genetic marker discovery and genotyping using next-generation sequencing. Nature Reviews Genetics, 12, 499–510.

Decroocq V, Fave MG, Hagen L, Bordenave L, Decroocq S (2003) Development and transferability of apricot and grape EST microsatellite markers across taxa. Theoretical and Applied Genetics, 106, 912–922.

Eaton DA (2014) PyRAD: assembly of de novo RADseq loci for phylogenetic analyses. Bioinformatics, 30, 1844–1849.

Etter PD, Bassham S, Hohenlohe PA, Johnson EA, Cresko WA (2012) SNP discovery and genotyping for evolutionary genetics using RAD sequencing. In: Molecular Methods for Evolutionary Genetics. Methods in Molecular Biology. pp. 157–178. Humana Press, Totowa, NJ.

Fernández ME, Goszczynski DE, Lirón JP et al. (2013) Comparison of the effectiveness of microsatellites and SNP panels for genetic identification, traceability and assessment of parentage in an inbred Angus herd. Genetics and Molecular Biology, 36, 185–191.

Galbusera P (2000) Cross-species amplification of microsatellite primers in passerine birds. Conservation Genetics, 1, 163–168.

Garvin MR, Saitoh K, Gharrett AJ (2010) Application of single nucleotide polymorphisms to non-model species: a technical review. Molecular Ecology Resources, 10, 915–934.

Glaubitz JC, Rhodes OE, Dewoody JA (2003) Prospects for inferring pairwise relationships with single nucleotide polymorphisms. Molecular Ecology, 12, 1039–1047.

Griffith SC, Owens IPF, Thuman KA (2002) Extra pair paternity in birds: a review of interspecific variation and adaptive function. Molecular Ecology, 11, 2195–2212.

Guichoux E, Lagache L, Wagner S et al. (2011) Current trends in microsatellite genotyping. Molecular Ecology Resources, 11, 591–611.

Gut IG (2001) Automation in genotyping of single nucleotide polymorphisms. Human Mutation, 17, 475–492.

Hadfield JD, Richardson DS, Burke T (2006) Towards unbiased parentage assignment: combining genetic, behavioural and spatial data in a Bayesian framework. Molecular Ecology, 15, 3715–3730.

Hauser L, Baird M, Hilborn R, Seeb LW, Seeb JE (2011) An empirical comparison of SNPs and microsatellites for parentage and kinship assignment in a wild sockeye salmon (Oncorhynchus nerka) population. Molecular Ecology Resources, 11, 150–161.

Hedgecock D, Li G, Hubert S, Bucklin K (2004) Widespread null alleles and poor crossspecies amplification of microsatellite DNA loci cloned from the Pacific oyster, Crassostrea gigas. Journal of Shellfish Research, 23, 379–385.

Heylar SJ, Hemmer Hansen J, Bekkevold D et al. (2011) Application of SNPs for population genetics of nonmodel organisms: new opportunities and challenges. Molecular Ecology Resources, 11, 123–136.

Hoffman JI, Amos W (2005) Microsatellite genotyping errors: detection approaches, common sources and consequences for paternal exclusion. Molecular Ecology, 14, 599–612.

Hughes C (1998) Integrating molecular techniques with field methods in studies of social behavior: a revolution results. Ecology, 79, 383–399.

Huisman, J (2017) Pedigree reconstruction from SNP data: parentage assignment, sibship clustering and beyond. Molecular Ecology Resources, 17, 1009–1024.

Kaiser SA, Taylor SA, Chen N et al. (2017) A comparative assessment of SNP and microsatellite markers for assigning parentage in a socially monogamous bird. Molecular Ecology Resources, 17, 183–193.

Karaket T, Poompuang S (2012) CERVUS vs. COLONY for successful parentage and sibship determinations in freshwater prawn Macrobrachium rosenbergii de Man. Aquaculture, 324, 307–311.

Kalinowski ST, Taper ML, Marshall TC (2007) Revising how the computer program CERVUS accommodates genotyping error increases success in paternity assignment. Molecular Ecology, 16, 1099–1106.

Kearse M, Moir R, Wilson A et al. (2012) Geneious Basic: An integrated and extendable desktop software platform for the organization and analysis of sequence data. Bioinformatics, 28, 1647–1649.

Kess T, Gross J, Harper, F, Boulding EG (2016) Low-cost ddRAD method of SNP discovery and genotyping applied to the periwinkle *Littorina saxatilis*. Journal of Molluscan Studies, 82, 104–109.

Li YC, Korol AB, Fahima T, Beiles A, Nevo E (2002) Microsatellites: genomic distribution, putative functions and mutational mechanisms: a review. Molecular Ecology, 11, 2453–2465.

Lischer HEL, Excoffier L (2012) PGDSpider: an automated data conversion tool for connecting population genetics and genomics programs. Bioinformatics, 28, 298–299.

Liu S, Palti Y, Gao G, Rexroad CE (2016) Development and validation of a SNP panel for parentage assignment in rainbow trout. Aquaculture, 452, 178–182.

Mastretta-Yanes A, Arrigo N, Alvarez N, Jorgensen TH, Piňero D, Emerson BC (2015) Restriction site-associated DNA sequencing, genotyping error estimation and de novo assembly optimization for population genetic inference. Molecular Ecology Resources, 15, 28–41.

Morin PA, Luikart G, Wayne RK (2004) SNPs in ecology, evolution and conservation. Trends in Ecology & Evolution, 19, 208–216.

Myers EM, Zamudio KR (2004) Multiple paternity in an aggregate breeding amphibian: the effect of reproductive skew on estimates of male reproductive success. Molecular Ecology, 13, 1951–1963.

Nali RC, Zamudio KR, Prado CPA (2014) Microsatellite markers for Bokermannohyla species (Anura, Hylidae) from the Brazilian Cerrado and Atlantic Forest domains. Amphibia-Reptilia, 35, 355–360.

Paris JR, Stevens JR, Catchen JM (2017) Lost in parameter space: a road map for Stacks. Methods in Ecology and Evolution. 8, 1360–1373.

Pemberton JM, Slate J, Bancroft DR, Barrett JA (1995) Nonamplifying alleles at microsatellite loci: a caution for parentage and population studies. Molecular Ecology, 4, 249–252.

Peterson BK, Weber JN, Kay EH, Fisher HS, Hoekstra HE (2012) Double digest RADseq: An inexpensive method for de novo SNP discovery and genotyping in model and non-model species (L Orlando, Ed,). PLoS ONE, 7, e37135.

Pew J, Muir PH, Wang J, Frasier TR (2015) related: an R package for analysing pairwise relatedness from codominant molecular markers. Molecular Ecology Resources, 15, 557–561.

Primmer CR, N Painter J, T Koskinen M, U Palo J, Merilä J (2005) Factors affecting avian cross-species microsatellite amplification. Journal of Avian Biology, 36, 348–360.

Puritz JB, Matz MV, Toonen RJ et al. (2014) Demystifying the RAD fad. Molecular Ecology, 23, 5937–5942.

Queller DC, Strassmann JE, Hughes CR (1993) Microsatellites and kinship. Trends in Ecology & Evolution, 8, 285–288.

R Core Team (2016) R: A Language and Environment for Statistical Computing. R Foundation for Statistical Computing, Vienna, Austria. http://www.R-project.org/.

Rohland N, Reich D (2012) Cost-effective, high-throughput DNA sequencing libraries for multiplexed target capture. Genome Research, 22, 939–946.

Ross KG (2001) Molecular ecology of social behaviour: analyses of breeding systems and genetic structure. Molecular Ecology, 10, 265–284.

Rowley I, Russell EM (1997) Fairy-wrens and Grasswrens: Maluridae. Oxford University Press.

Saunders IW, Brohede J, Hannan GN (2007) Estimating genotyping error rates from Mendelian errors in SNP array genotypes and their impact on inference. Genomics, 90, 291–296.

Schodde, R (1982). The fairy-wrens (ed. Bass T). The Craftsman Press. Victoria, Australia.

Seeb JE, Carvalho G, Hauser L et al. (2011) Single-nucleotide polymorphism (SNP) discovery and applications of SNP genotyping in nonmodel organisms. Molecular Ecology Resources, 11, 1–8.

Selkoe KA, Toonen RJ (2006) Microsatellites for ecologists: a practical guide to using and evaluating microsatellite markers. Ecology Letters, 9, 615–629.

Shafer A, Peart CR, Tusso S, Maayan I, Brelsford A, Wheat CW, Wolf JB (2016) Bioinformatic processing of RAD-seq data dramatically impacts downstream population genetic inference. Methods in Ecology and Evolution, 8, 907–917.

Slate J, Gratten J, Beraldi D et al. (2010) Gene mapping in the wild with SNPs: guidelines and future directions. Genetica, 136, 97–107.

Solomon NG, Keane B, Knoch LR (2004) Multiple paternity in socially monogamous prairie voles (Microtus ochrogaster). Canadian Journal of Zoology, 82, 1667–1671.

Strucken EM, Lee SH, Lee HK et al. (2016) How many markers are enough? Factors influencing parentage testing in different livestock populations. Journal of Animal Breeding and Genetics, 133, 13–23.

Syvänen A-C (2001) Accessing genetic variation: genotyping single nucleotide polymorphisms. Nature Reviews Genetics, 2, 930–942.

Tokarska M, Marshall T, Kowalczyk R et al. (2009) Effectiveness of microsatellite and SNP markers for parentage and identity analysis in species with low genetic diversity: the case of European bison. Heredity, 103, 326–332.

Tortereau F, Moreno CR, Tosser-Klopp G, Servin B, Raoul J (2017) Development of a SNP panel dedicated to parentage assignment in French sheep populations. BMC Genetics, 18, 50.

Wang J (2002) An estimator for pairwise relatedness using molecular markers. Genetics, 160, 1203–1215.

Wang J (2007) Triadic IBD coefficients and applications to estimating pairwise relatedness. Genetical Research, 89, 135–153.

Webster MS, Reichart L (2005) Use of microsatellites for parentage and kinship analyses in animals. Methods in Enzymology, 395, 222–238.

Weinman LR, Solomon JW, Rubenstein DR (2015) A comparison of single nucleotide polymorphism and microsatellite markers for analysis of parentage and kinship in a cooperatively breeding bird. Molecular Ecology Resources, 15, 502–511.

Westneat DF, Sherman PW, Morton ML (1990) The ecology and evolution of extra-pair copulations in birds. Current Ornithology, 7, 331–370.

White PS, Densmore LD (1992) Mitochondrial DNA isolation. In: Molecular Genetic Analysis of Populations: A Practical Approach (ed. Hoelzel AR), pp. 50–51. Oxford University Press, New York, NY.

Wigginton, JE, Cutler DJ, Abecasis GR (2005) A note on exact tests of Hardy-Weinberg equilibrium. American Journal of Human Genetics, 76, 887–893.

Willing E-M, Hoffmann M, Klein JD, Weigel D, Dreyer C (2011) Paired-end RAD-seq for de novo assembly and marker design without available reference. Bioinformatics, 27, 2187–2193.

